# Differential Striatal Axonal Arborizations of the Intratelencephalic and Pyramidal-Tract Neurons: Analysis of the Data in the MouseLight Database

**DOI:** 10.1101/752824

**Authors:** Kenji Morita, Sanghun Im, Yasuo Kawaguchi

## Abstract

There exist two major types of striatum-targeting neocortical neurons, specifically, intratelencephalic (IT) neurons and pyramidal-tract (PT) neurons. Regarding their striatal projections, it was once suggested that IT axons are extended whereas PT axons are primarily focal. However, subsequent study with an increased number of well-stained extended axons concluded that such an apparent distinction was spurious due to limited sample size. Recent work using genetically labeled neurons reintroduced the differential spatial extent of the striatal projections of IT and PT neurons through population-level analyses, complemented by observations of single axons. However, quantitative analysis of a large number of axons remained to be conducted. We analyzed the data of axonal end-points of 161 IT neurons and 33 PT neurons in the MouseLight database (http://ml-neuronbrowser.janelia.org/). The number of axonal end-points in the ipsilateral striatum exhibits roughly monotonically decreasing distributions in both neuron types. However, the proportion of neurons having >50 ipsilateral end-points is larger in IT neurons than in PT neurons. Moreover, distinguishing IT subpopulations in the secondary motor area (MOs), layer 5 neurons and bilateral striatum-targeting layer 2/3 neurons, but not contralateral striatum-non-targeting layer 2/3 neurons, have a larger number of ipsilateral end-points than MOs PT neurons. We also found that IT ipsilateral striatal axonal end-points are on average more widely distributed than PT end-points, especially in the medial-lateral direction. These results indicate that IT and PT striatal axons differ in the frequency of having numerous end-points and the spatial extent of end-points while there are wide varieties within each neuron type.

## Introduction

There exist two major types of striatum targeting neurons in the neocortex, specifically, intratelencephalic (IT) neurons, which project only to telencephalic regions, and pyramidal-tract (PT) neurons, which project out of telencephalon (Wilson, 1986, 1987; Cowan and Wilson, 1994; Levesque et al., 1996; Parent and Parent, 2006; Reiner et al., 2010; Shepherd, 2013). These neuron types commonly exist in neocortical areas including the motor cortices, and have distinct, albeit overlapped, layer distributions. They also have differential dendritic morphology and intracortical connectivity (Morishima and Kawaguchi, 2006; Brown and Hestrin, 2009; Morishima et al., 2011; Kiritani et al., 2012), as well as gene expression (Arlotta et al., 2005; Molyneaux et al., 2009; Tasic et al., 2018). Regarding the striatal projections of IT and PT neurons, contralateral projections arise only from IT neurons. Moreover, it was once suggested that IT axonal arborizations are extended whereas PT axons are primarily focal, based on intracellular labeling of IT and PT neurons identified by antidromic activation from the contralateral striatum or the medullary pyramid, respectively (Cowan and Wilson, 1994). However, subsequent study from the same laboratory examined an increased number of well-stained extended axons (10 IT neurons and 6 PT neurons), and concluded that such an apparent difference in the axonal morphology was spurious due to limited sample size (Zheng and Wilson, 2002).

Recent work (Hooks et al., 2018) systematically examined the striatal projections of IT and PT neurons by injecting Cre-dependent fluorescent reporters into various cortical sites in mouse lines specifically expressing Cre in either IT or PT neurons (Gerfen et al., 2013). Along with revealing the neuron type- and cortical area-dependent topographic precision, which was the main focus of the study, the authors have shown that cortical injection of Cre-dependent reporter in the PT-Cre mouse line caused fewer striatal voxels with suprathreshold fluorescence intensity (Hooks et al., 2018), indicating that PT projections are spatially more limited at the population level. Moreover, they complemented their argument by observing axonal arborizations of IT and PT neurons collected through the MouseLight project at Janelia Research Campus, which performed whole brain reconstructions (Economo et al., 2016). Based on the observation, they mentioned that IT axons were more extensive and PT axons were more focal, and they also referred to the previous study (Cowan and Wilson, 1994) (but not (Zheng and Wilson, 2002)). However, quantitative analysis of a large number of axonal morphology data was not performed.

The MouseLight project has now developed the MouseLight database (http://ml-neuronbrowser.janelia.org/), which contains reconstructed morphology data of about 1000 neurons and is open to public (Economo et al., 2019; Winnubst et al., 2019). The article introducing this database (Winnubst et al., 2019) performed several analyses including those for IT and PT neurons, which revealed great diversity of IT neuronal projection patterns and also PT neuronal subtypes projecting to distinct targets. However, quantitative comparison of intra-striatal axons between IT and PT neurons was not reported. Because this long-standing issue is critical in elucidating the functions of corticostriatal circuits, we addressed it by using the MouseLight database.

## Methods

### Identification of PT and IT neurons in the MouseLight database

The source of the data used in this article is the MouseLight project at Janelia, and the DOIs of data entities are listed in Table 1. We searched entities of pyramidal-tract (PT) -type corticostriatal neurons in the Neuron Browser of the MouseLight database (http://ml-neuronbrowser.janelia.org/) by setting three filters: (1) soma is located in the cerebral cortex (“Cerebral cortex” in the search box), (2) axonal end-point exists in the striatum (specifically, “Striatum”, “Striatum dorsal region”, “Striatum ventral region”, “Caudoputamen” or “Nucleus accumbens”; threshold: “any”), and (3) axon exists in the pons (specifically, “Pons”, “Pons, sensory related”, “Pons, motor related”, or “Pons, behavioral state related”; threshold: “any”). As a result, we found 36 entities of neocortical neurons with all the structures; 3 entities with “axon only” and a subiculum neuron were omitted from our analyses. In these 36 entities, 33 entities have soma in layer 5 whereas 2 have soma in layer 2/3 and 1 lacks description of soma layer. Since PT neurons have been described to have soma in layer 5 in previous studies, we included the 33 neurons with soma in layer 5 into analyses as PT neurons.

**Table 1.** List of ID number and DOI of each neuron data entity used in this article. The source of the data is the MouseLight project at Janelia (http://ml-neuronbrowser.janelia.org/).

| ID | DOI |
| --- | --- |
| **PT in the primary motor area (MOp)** |  |
| AA0132 | 10.25378/janelia.5527270 |
| AA0135 | 10.25378/janelia.5527279 |
| AA0261 | 10.25378/janelia.5527717 |
| AA0584 | 10.25378/janelia.7649918 |
| AA0587 | 10.25378/janelia.7649927 |
| AA0617 | 10.25378/janelia.7655711 |
| AA0923 | 10.25378/janelia.7803737 |
| AA0927 | 10.25378/janelia.7803770 |
| **PT in the secondary motor area (MOs)** |  |
| AA0011 | 10.25378/janelia.5521615 |
| AA0012 | 10.25378/janelia.5521618 |
| AA0122 | 10.25378/janelia.5527240 |
| AA0180 | 10.25378/janelia.5527441 |
| AA0181 | 10.25378/janelia.5527444 |
| AA0182 | 10.25378/janelia.5527447 |
| AA0245 | 10.25378/janelia.5527657 |
| AA0250 | 10.25378/janelia.5527678 |
| AA0415 | 10.25378/janelia.7614212 |
| AA0583 | 10.25378/janelia.7649900 |
| AA0726 | 10.25378/janelia.7707194 |
| AA0764 | 10.25378/janelia.7710065 |
| AA0780 | 10.25378/janelia.7739285 |
| AA0788 | 10.25378/janelia.7739369 |
| AA0791 | 10.25378/janelia.7739399 |
| AA0792 | 10.25378/janelia.7739402 |
| **PT in other areas** |  |
| AA0001 | 10.25378/janelia.5520037 |
| AA0066 | 10.25378/janelia.5521807 |
| AA0119 | 10.25378/janelia.5526736 |
| AA0121 | 10.25378/janelia.5527237 |
| AA0796 | 10.25378/janelia.7739525 |
| AA0941 | 10.25378/janelia.7804004 |
| AA0944 | 10.25378/janelia.7804028 |
| AA0945 | 10.25378/janelia.7804034 |
| AA0956 | 10.25378/janelia.7804088 |
| **IT in MOp** |  |
| AA0002 | 10.25378/janelia.5520049 |
| AA0004 | 10.25378/janelia.5520205 |
| AA0010 | 10.25378/janelia.5521600 |
| AA0034 | 10.25378/janelia.5521684 |
| AA0035 | 10.25378/janelia.5521690 |
| AA0042 | 10.25378/janelia.5521714 |
| AA0062 | 10.25378/janelia.5521789 |
| AA0064 | 10.25378/janelia.5521798 |
| AA0065 | 10.25378/janelia.5521801 |
| AA0099 | 10.25378/janelia.5526676 |
| AA0102 | 10.25378/janelia.5526685 |
| AA0107 | 10.25378/janelia.5526700 |
| AA0108 | 10.25378/janelia.5526703 |
| AA0130 | 10.25378/janelia.5527264 |
| AA0140 | 10.25378/janelia.5527297 |
| AA0184 | 10.25378/janelia.5527453 |
| AA0271 | 10.25378/janelia.5527762 |
| AA0272 | 10.25378/janelia.5527765 |
| AA0289 | 10.25378/janelia.5527822 |
| AA0401 | 10.25378/janelia.7614125 |
| AA0404 | 10.25378/janelia.7614134 |
| AA0408 | 10.25378/janelia.7614173 |
| AA0440 | 10.25378/janelia.7614341 |
| AA0442 | 10.25378/janelia.7614368 |
| AA0541 | 10.25378/janelia.7640162 |
| AA0543 | 10.25378/janelia.7640168 |
| AA0578 | 10.25378/janelia.7649858 |
| AA0582 | 10.25378/janelia.7649873 |
| AA0588 | 10.25378/janelia.7649933 |
| AA0592 | 10.25378/janelia.7649948 |
| AA0600 | 10.25378/janelia.7650029 |
| AA0618 | 10.25378/janelia.7655729 |
| AA0622 | 10.25378/janelia.7655753 |
| AA0627 | 10.25378/janelia.7655771 |
| AA0656 | 10.25378/janelia.7658219 |
| AA0662 | 10.25378/janelia.7658243 |
| AA0663 | 10.25378/janelia.7658246 |
| AA0674 | 10.25378/janelia.7704212 |
| AA0739 | 10.25378/janelia.7707326 |
| AA0741 | 10.25378/janelia.7707338 |
| AA0745 | 10.25378/janelia.7707359 |
| AA0876 | 10.25378/janelia.7742576 |
| AA0884 | 10.25378/janelia.7742789 |
| AA0906 | 10.25378/janelia.7780859 |
| IT in MOs layer 2/3, Contralateral-Striatum-non-targeting |  |
| AA0014 | 10.25378/janelia.5521624 |
| AA0116 | 10.25378/janelia.5526727 |
| AA0118 | 10.25378/janelia.5526733 |
| AA0237 | 10.25378/janelia.5527630 |
| AA0238 | 10.25378/janelia.5527633 |
| AA0241 | 10.25378/janelia.5527645 |
| AA0418 | 10.25378/janelia.7614221 |
| AA0426 | 10.25378/janelia.7614251 |
| AA0446 | 10.25378/janelia.7614707 |
| AA0471 | 10.25378/janelia.7615952 |
| AA0474 | 10.25378/janelia.7615964 |
| AA0738 | 10.25378/janelia.7707323 |
| AA0782 | 10.25378/janelia.7739303 |
| AA0793 | 10.25378/janelia.7739510 |
| AA0802 | 10.25378/janelia.7739555 |
| AA0803 | 10.25378/janelia.7739558 |
| AA0915 | 10.25378/janelia.7780892 |
| IT in MOs layer 2/3, Contralateral-Striatum-targeting |  |
| AA0232 | 10.25378/janelia.5527609 |
| AA0327 | 10.25378/janelia.7613498 |
| AA0328 | 10.25378/janelia.7613507 |
| AA0329 | 10.25378/janelia.7613525 |
| AA0407 | 10.25378/janelia.7614158 |
| AA0409 | 10.25378/janelia.7614182 |
| AA0416 | 10.25378/janelia.7614215 |
| AA0419 | 10.25378/janelia.7614227 |
| AA0439 | 10.25378/janelia.7614329 |
| AA0450 | 10.25378/janelia.7614965 |
| AA0467 | 10.25378/janelia.7615901 |
| AA0470 | 10.25378/janelia.7615940 |
| AA0773 | 10.25378/janelia.7710113 |
| AA0865 | 10.25378/janelia.7740089 |
| AA0866 | 10.25378/janelia.7740092 |
| AA0873 | 10.25378/janelia.7742567 |
| AA0883 | 10.25378/janelia.7742786 |
| AA0897 | 10.25378/janelia.7780811 |
| IT in MOs layer 5 |  |
| AA0059 | 10.25378/janelia.5521780 |
| AA0190 | 10.25378/janelia.5527474 |
| AA0230 | 10.25378/janelia.5527603 |
| AA0233 | 10.25378/janelia.5527612 |
| AA0236 | 10.25378/janelia.5527621 |
| AA0265 | 10.25378/janelia.5527738 |
| AA0267 | 10.25378/janelia.5527747 |
| AA0269 | 10.25378/janelia.5527753 |
| AA0274 | 10.25378/janelia.5527774 |
| AA0279 | 10.25378/janelia.5527792 |
| AA0281 | 10.25378/janelia.5527798 |
| AA0285 | 10.25378/janelia.5527810 |
| AA0300 | 10.25378/janelia.5527855 |
| AA0324 | 10.25378/janelia.7613486 |
| AA0332 | 10.25378/janelia.7613684 |
| AA0397 | 10.25378/janelia.7614113 |
| AA0400 | 10.25378/janelia.7614122 |
| AA0412 | 10.25378/janelia.7614200 |
| AA0421 | 10.25378/janelia.7614233 |
| AA0422 | 10.25378/janelia.7614236 |
| AA0441 | 10.25378/janelia.7614356 |
| AA0452 | 10.25378/janelia.7615274 |
| AA0460 | 10.25378/janelia.7615835 |
| AA0465 | 10.25378/janelia.7615889 |
| AA0466 | 10.25378/janelia.7615895 |
| AA0473 | 10.25378/janelia.7615961 |
| AA0534 | 10.25378/janelia.7640063 |
| AA0575 | 10.25378/janelia.7649846 |
| AA0602 | 10.25378/janelia.7650038 |
| AA0632 | 10.25378/janelia.7658054 |
| AA0646 | 10.25378/janelia.7658126 |
| AA0734 | 10.25378/janelia.7707296 |
| AA0735 | 10.25378/janelia.7707302 |
| AA0746 | 10.25378/janelia.7707365 |
| AA0749 | 10.25378/janelia.7707374 |
| AA0767 | 10.25378/janelia.7710077 |
| AA0798 | 10.25378/janelia.7739537 |
| AA0841 | 10.25378/janelia.7739909 |
| AA0842 | 10.25378/janelia.7739915 |
| AA0853 | 10.25378/janelia.7739954 |
| AA0858 | 10.25378/janelia.7740017 |
| AA0887 | 10.25378/janelia.7742807 |
| AA0905 | 10.25378/janelia.7780856 |
| IT in MOs other layer or without layer description |  |
| AA0100 | 10.25378/janelia.5526679 |
| AA0106 | 10.25378/janelia.5526697 |
| AA0113 | 10.25378/janelia.5526718 |
| AA0243 | 10.25378/janelia.5527651 |
| AA0288 | 10.25378/janelia.5527819 |
| AA0320 | 10.25378/janelia.7613465 |
| AA0333 | 10.25378/janelia.7613693 |
| AA0396 | 10.25378/janelia.7614107 |
| AA0402 | 10.25378/janelia.7614128 |
| AA0461 | 10.25378/janelia.7615838 |
| AA0549 | 10.25378/janelia.7640201 |
| AA0594 | 10.25378/janelia.7649966 |
| AA0645 | 10.25378/janelia.7658114 |
| AA0653 | 10.25378/janelia.7658153 |
| AA0655 | 10.25378/janelia.7658210 |
| AA0742 | 10.25378/janelia.7707347 |
| AA0743 | 10.25378/janelia.7707350 |
| AA0744 | 10.25378/janelia.7707356 |
| AA0748 | 10.25378/janelia.7707371 |
| AA0790 | 10.25378/janelia.7739384 |
| AA0880 | 10.25378/janelia.7742777 |
| AA0889 | 10.25378/janelia.7742816 |
| AA0911 | 10.25378/janelia.7780874 |
| IT in other areas |  |
| AA0008 | 10.25378/janelia.5520451 |
| AA0098 | 10.25378/janelia.5526673 |
| AA0120 | 10.25378/janelia.5527234 |
| AA0319 | 10.25378/janelia.7613459 |
| AA0393 | 10.25378/janelia.7614098 |
| AA0403 | 10.25378/janelia.7614131 |
| AA0417 | 10.25378/janelia.7614218 |
| AA0425 | 10.25378/janelia.7614245 |
| AA0427 | 10.25378/janelia.7614254 |
| AA0590 | 10.25378/janelia.7649942 |
| AA0603 | 10.25378/janelia.7650041 |
| AA0679 | 10.25378/janelia.7704230 |
| AA0795 | 10.25378/janelia.7739519 |
| AA0800 | 10.25378/janelia.7739543 |
| AA0801 | 10.25378/janelia.7739549 |
| AA0872 | 10.25378/janelia.7742564 |
| Corticostriatal neurons that we did not include into our analyses as IT or PT neurons |  |
| AA0096 | 10.25378/janelia.5526667 |
| AA0105 | 10.25378/janelia.5526694 |
| AA0231 | 10.25378/janelia.5527606 |
| AA0284 | 10.25378/janelia.5527807 |
| AA0395 | 10.25378/janelia.7614104 |
| AA0413 | 10.25378/janelia.7614203 |
| AA0445 | 10.25378/janelia.7614596 |
| AA0457 | 10.25378/janelia.7615589 |
| AA0469 | 10.25378/janelia.7615934 |
| AA0492 | 10.25378/janelia.7616045 |
| AA0636 | 10.25378/janelia.7658072 |
| AA0641 | 10.25378/janelia.7658087 |
| AA0642 | 10.25378/janelia.7658090 |
| AA0644 | 10.25378/janelia.7658108 |
| AA0647 | 10.25378/janelia.7658135 |
| AA0668 | 10.25378/janelia.7658261 |
| AA0671 | 10.25378/janelia.7704200 |
| AA0747 | 10.25378/janelia.7707368 |
| AA0765 | 10.25378/janelia.7710068 |
| AA0784 | 10.25378/janelia.7739321 |
| AA0807 | 10.25378/janelia.7739642 |
| AA0840 | 10.25378/janelia.7739903 |
| AA0870 | 10.25378/janelia.7742552 |
| AA0900 | 10.25378/janelia.7780826 |
| AA0914 | 10.25378/janelia.7780883 |
| AA0919 | 10.25378/janelia.7780907 |

Next we searched entities of intratelencephalic (IT) -type corticostriatal neurons. A potential strategy was to search neurons having axons in the contralateral striatum, since previous studies have described that only IT neurons, but not PT neurons, can project to the contralateral striatum (Miller, 1975; Catsman-Berrevoets et al., 1980; Wilson, 1986). However, because IT neurons do not necessarily project to the contralateral striatum and also because practically we could not find a way to specify the contralateral striatum in the Neuron Browser, we took a different strategy. Specifically, we conducted a separate search of the Neuron Browser by setting only filters (1) and (2) mentioned above, omitting filter (3) (axon in pons), and manually excluded entities that were also found in the search with filter (3) so as to obtain candidates of IT neurons. This yielded 187 entities of neocortical neurons with all the structures; neurons with soma not located in the neocortex (but in the hippocampus or hippocampal formation in most cases) were also manually excluded.

In order to check if these 187 neurons satisfy the definition of IT neurons, i.e., axon projections only within the telencephalon, we examined json files of these entities downloaded from the MouseLight database and checked if axon exists in “allenId”s that are considered to be outside of the telencephalon (but omitting some “allenId”s corresponding to tract, bundle, or ventricle, parts of which could potentially be intratelencephalic, such as the corticospinal tract). As a result, 26 (out of 187) entities were found to have axon in non-IT regions (Table 2). Of these 26 entities, 11 are neurons with a substantial portion of axons (> 10% of axon entities in json file) in non-IT regions, such as thalamus, midbrain, or hypothalamus, and would thus be inappropriate to be labeled as IT neurons. Among them, AA0919 and AA0644 having soma in layer 5 could potentially be PT neurons, although we did not include them into our analyses as PT neurons. Other than these and a neuron having soma in layer 2/3, 8 out of the 11 neurons having soma in layer 6a and targeting thalamus would be corticothalamic neurons. Layer 6 corticothalamic neurons are distinct from PT neurons, and they together constitute extratelencephalic neurons (Baker et al., 2018). Layer 6 striatum-targeting corticothalamic neurons can thus be regarded as a third type of corticostriatal neurons (i.e., other than IT and PT neurons), but here we did not further analyze them. The remaining 15 entities are neurons whose non-IT axon projections are limited (< 2.2% of axon entities in json file), and it might be good to classify them together with properly IT neurons, although we did not do so.

**Table 2.**
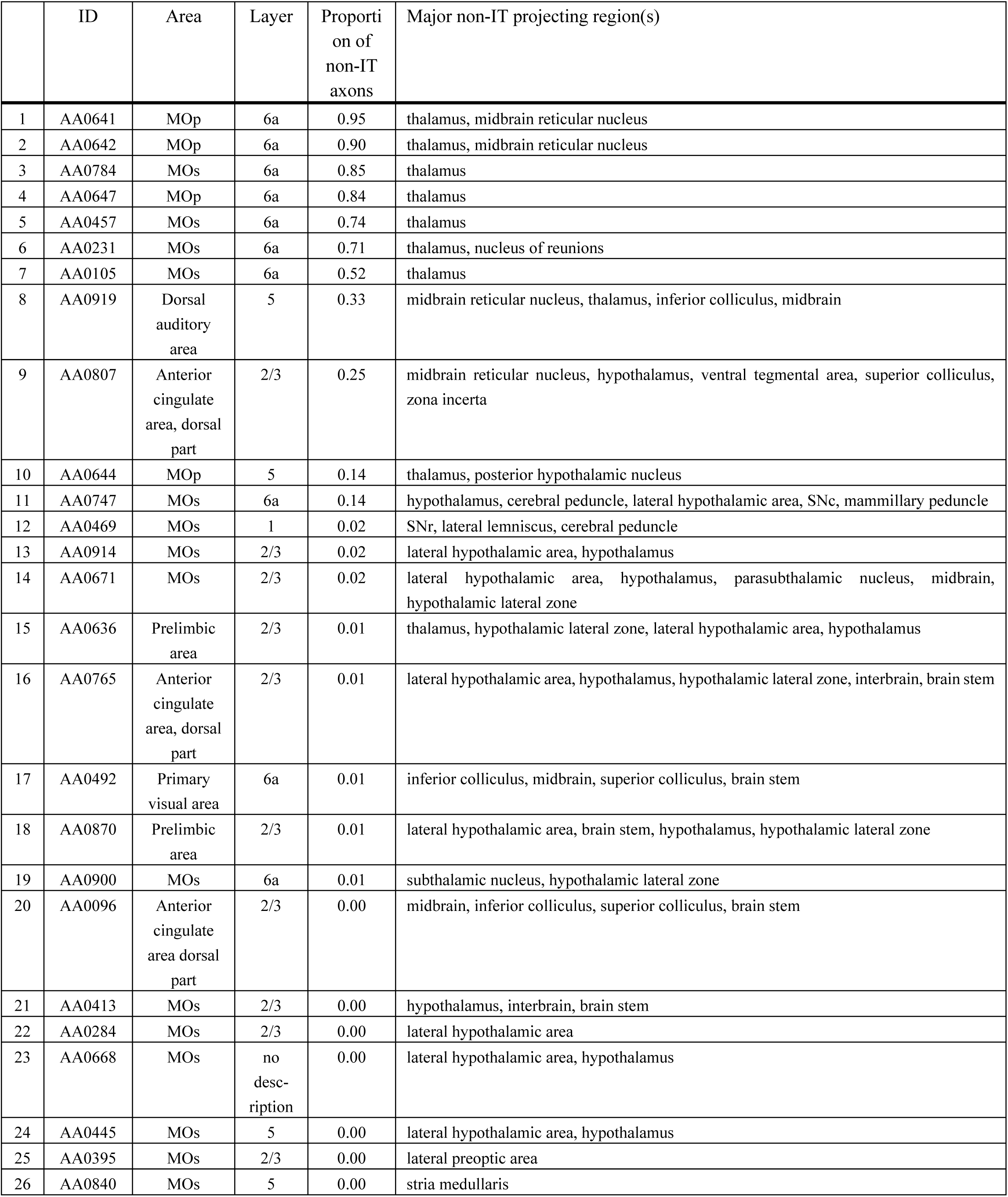
List of entities of corticostriatal neurons in the MouseLight database that we did not include into our analyses as IT or PT neurons.

In the end, we analyzed 161 (= 187 - 26) entities as IT neurons. Of these, 44 neurons are in the primary motor area (3 in layer 1 (according to the annotation in the MouseLight database), 6 in layer 2/3, 23 in layer 5, and 12 in layer 6a), 101 neurons are in the secondary motor area (4 in layer 1, 35 in layer 2/3, 43 in layer 5, 17 in layer 6a, and 2 without description of layer), and the remaining 16 neurons are in other neocortical areas. Also, we analyzed 33 entities as PT neurons (all with soma in layer 5 as mentioned above). Of these, 8, 16, and 9 neurons are in the primary motor area, secondary motor area, and other neocortical areas, respectively. Figure 1 shows the axon morphology of examples of PT and IT neurons. (Note: although we did not record the numbers of search results at our original searches in the MouseLight database, we later realized that additionally specifying “Fundus of striatum” and “Olfactory tubercle” for filter (2), or specifying “Striatum” only for filter (2) and “Pons” only for filter (3), does not change the number of search results).

**Figure 1.**
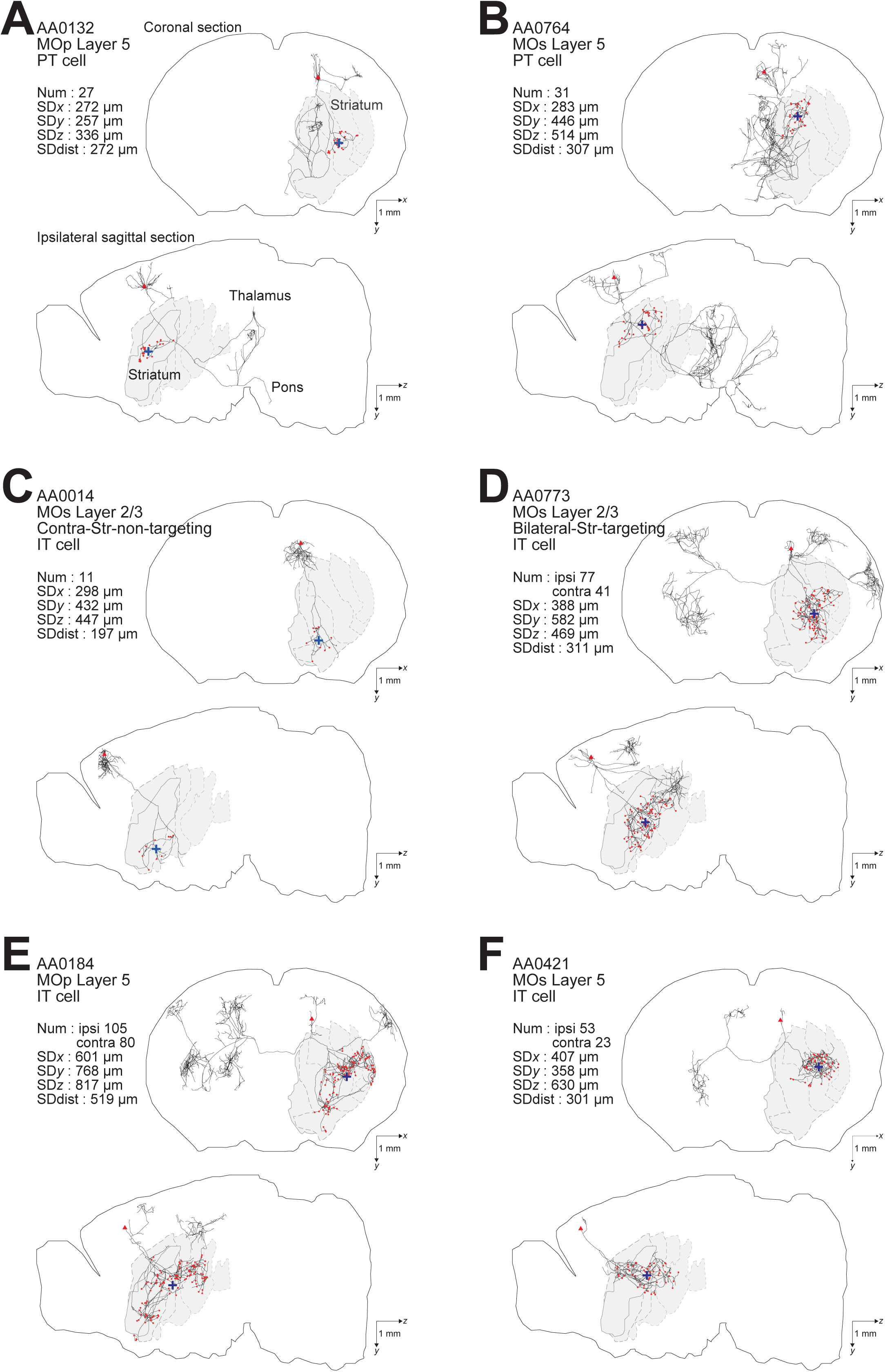
Examples of axonal arborizations of PT and IT neurons, drawn by using the data downloaded from the MouseLight database (http://ml-neuronbrowser.janelia.org/). Coronal (*x*-*y* plane) and sagittal (*y*-*z* plane) sections are shown for individual cells. The red points indicate the axonal end-points ipsilateral to the soma, and the blue cross indicates the center of those end-points (i.e., the point whose coordinates are the means of the *x, y*, and *z* coordinates of individual end-points). The small red triangle indicates the soma. The text in the top-left of each panel describes the following information: ID number in the MouseLight database; Cortical area of soma (primary motor area (MOp) or secondary motor area (MOs)); Layer of soma, Neuron type (PT or IT); Num: number of axonal end-points in the ipsilateral striatum (A-C) or in the bilateral striatum (D-F); SD*x*,SD*y*,SD*z*: the standard deviation (SD) of *x, y*, and *z* coordinates of the ipsilateral intra-striatal end-points (in [μm]); and SDdist: SD of the distances of ipsilateral intra-striatal end-points from the center of those end-points (in [μm]). The contours of striatum were referred from Allen 3-D annotation (Kuan et al., 2015) and filled in gray.

### Analysis of axonal end-points in the striatum

We analyzed json files of the identified IT and PT neurons downloaded from the MouseLight database to identify axonal end-points in the striatum, i.e., axon samples (“structureIdentifier”: 2) that have “allenId” of 477 (Striatum), 485 (Striatum dorsal region), 493 (Striatum ventral region), 672 (Caudoputamen), 56 (Nucleus accumbens), 754 (Olfactory tubercle), or 998 (Fundus of striatum) and are not a parent of other axon samples (i.e., whose “sampleNumber” does not appear as “parentNumber” of other axon samples). We classified the identified axonal end-points into either ipsi- or contra-lateral axonal end-points by examining whether the *x* coordinate (right-left position) is at the same side as the soma with respect to *x*=5500, which appeared to be near the midline when we plotted the distribution of *x* coordinates of striatal axonal end-points in an example neuron. Later we realized that *x*=5700 would actually be around the midline, but we have confirmed that the number of ipsi/contralateral intra-striatal axonal end-points and their coordinates for all the IT and PT neurons do not change when *x*=5700, instead of *x*=5500, is used in the extraction.

We did analyses with MATLAB (MathWorks Inc.), using custom-made codes and the codes for statistical analyses formerly in http://rnpsychology.org/ (by Ryosuke Niimi), and R (https://www.r-project.org/). Rounding errors were introduced when data were moved from MATLAB to CSV files using csvwrite.m for analyses using R.

## Results

### Number of intra-striatal axonal end-points

We examined how the number of intra-striatal axonal end-points is distributed across neuron types as well as across individual neurons. As shown in Fig. 2Aa, the number turned out to be widely distributed across individual neurons in both PT and IT neural populations and for both ipsi- and contra-lateral axonal end-points in the case of IT neurons (no PT neuron has axonal end-point in the contra-lateral striatum, consistent with previous studies). In all the cases, the distributions are roughly monotonically decreasing, i.e., neurons with ≤20 axonal end-points are most frequent, while there are also neuron(s) having ≥100 axonal end-points. Comparing the ipsi- and contra-lateral axonal end-points in IT neurons, there tend to be more ipsilateral end-points than contra-lateral end-points: ipsi-points outnumber contra-points in 122 (out of 161) neurons, including 42 neurons without contra-points, whereas contra-points outnumber ipsi-points in 38 neurons, including 16 neurons without ipsi-points (the remaining 1 neuron has the same numbers of ipsi- and contra-points).

**Figure 2.**
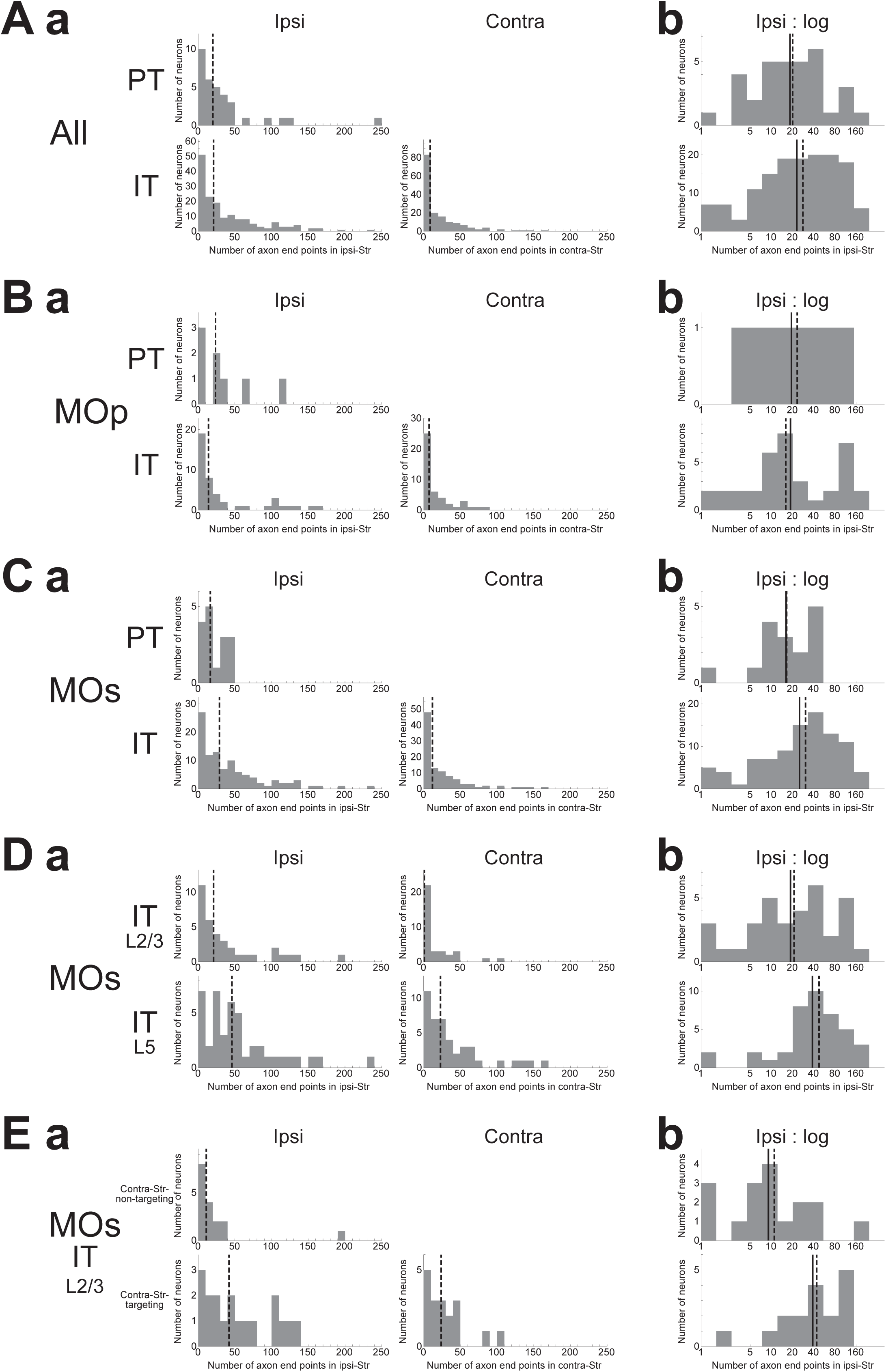
Distributions of the number of intra-striatal axonal end-points. **(A) (a)** Distributions for all the striatum-targeting PT and IT neurons (33 and 161 neurons, respectively) that we identified in the MouseLight database. IT neurons having no ipsi-(16) or contra-(42) lateral axonal end-point are included in the histograms. The dashed lines indicate the medians. **(b)** Distributions of the logarithm of the number of ipsilateral intra-striatal axonal end-points for the striatum-targeting 33 PT and 145 IT neurons that have at least one ipsilateral end-point; i.e., IT neurons having no ipsilateral axonal end-point (16 neurons) are excluded from the histogram (the same is applied also to the following histograms in (b)). The solid lines and the dashed lines indicate the means and the medians, respectively; the same is applied also to the following figures. **(B**,**C)** Results for the neurons whose somata are located in the primary motor area ((a) 8 PT and 44 IT neurons (including 7 w/o ipsi-point and 10 w/o contra-point), (b) 8 PT and 37 IT neurons) (B) or the secondary motor area (MOs) ((a) 16 PT and 101 IT neurons (including 7 w/o ipsi-point and 27 w/o contra-point), (b) 16 PT and 94 IT neurons) (C). **(D)** Results for MOs layer 2/3 IT neurons ((a) 35 neurons including 1 w/o ipsi-point and 17 w/o contra-point, (b) 34 neurons) and MOs layer 5 IT neurons ((a) 43 neurons including 3 w/o ipsi-point and 4 w/o contra-point, (b) 40 neurons). **(E)** Results for MOs layer 2/3 IT neurons that do not target contralateral striatum (contra-Str) (17 neurons) and those that target contra-Str ((a) 18 neurons including 1 w/o ipsi-point, (b) 17 neurons).

Comparing the ipsilateral axonal end-points between PT and IT neurons, the proportion of neurons having a large number (>50) of ipsi-points within those having at least one ipsi-point is larger in IT neurons (48/145=0.33) than in PT neurons (5/33=0.15) (*χ*^2^ test, *p*=0.042, *φ*=0.15). Nonetheless, the intra-neuron-type variabilities across individual neurons looks more prominent than the interneuron type variability. Indeed, even the distributions of the logarithm of the number of ipsilateral axonal end-points, excluding the neurons having no ipsilateral end-point (this is also applied to all the following analyses dealing with the logarithm of the number of end-points so as to avoid “log 0”), are considerably overlapped between PT and IT neurons (Fig. 2Ab) (Welch’s *t* test, *p*=0.36). Figure 2B and C are the results of analyses limited to neurons in the primary motor area (MOp) or secondary motor area (MOs), respectively, showing similar tendencies to the results for all the neurons.

We also analyzed if MOs IT neurons in layer 2/3 and those in layer 5 differ in the number of intra-striatal axonal end-points (Fig. 2D). It turned out that whereas most layer 5 MOs IT neurons (39 out of 43) have at least one axonal end-point in the contralateral striatum, only about a half of layer 2/3 MOs IT neurons (18 out of 35) have contralateral striatal end-point(s). There is also a trend that the proportion of neurons having a large number (>50) of ipsilateral end-points within those having at least one ipsi-point tends to be larger in layer 5 MOs IT neurons (18/40=0.45) than in layer 2/3 neurons (9/34=0.26) (*χ*^2^ test, *p*=0.099, *φ*=0.19). Moreover, the distributions of the logarithm of the number of ipsilateral axonal end-points, excluding the neurons having no ipsilateral end-point, differ between layer 2/3 and layer 5 MOs IT neurons (Welch’s *t*-test, *p*=0.024, *d*=0.55), with the layer 5 neurons on average having a larger number of ipsilateral end-points than the layer 2/3 neurons as apparent in Fig. 2Db. The distributions also differ between MOs layer 5 IT neurons and MOs PT neurons (*p*=0.0097, *d*=0.76) but not between MOs layer 2/3 IT neurons and MOs PT neurons (*p*=0.68).

As mentioned above, most of MOs layer 5 IT neurons (39/43) project to contralateral striatum (contra-Str), whereas MOs layer 2/3 IT neurons are almost bisected into those targeting contra-Str (18/35) and those not targeting (17/35). Except for a contra-Str-targeting neurons that does not target ipsilateral striatum, the remaining bilateral-Str-targeting neurons (17) on average have a larger number of ipsilateral end-points than the contra-Str-non-targeting neurons (Fig. 2E), with the distributions of the logarithm of the number of end-points significantly different (Welch’s *t*-test, *p*=0.0022, *d*=1.15). The distribution for bilateral-Str-targeting MOs layer 2/3 IT neurons does not differ from that for MOs layer 5 IT neurons (*p*=0.97) but differs from that for MOs PT neurons (*p*=0.024, *d*=0.82). On the contrary, the distribution for contra-Str-non-targeting MOs layer 2/3 IT neurons differs from that for MOs layer 5 IT neurons (*p*=0.0010, *d*=1.16) but hardly differs from that for MOs PT neurons (*p*=0.19).

### Spatial distribution of intra-striatal axonal end-points

We also examined how the ipsilateral intra-striatal axonal end-points are spatially distributed. Specifically, we calculated the standard deviation (SD) of *x, y*, and *z* coordinates (corresponding to the medial-lateral (or right-left), dorsal-ventral (or top-bottom), and anterior-posterior directions, respectively) of the ipsilateral end-point(s) for each neuron, excluding the neurons having no ipsilateral end-point (this is also applied to all the following analyses dealing with the SD of the coordinates). As shown in Fig. 3A, the SD is distributed from 0 to several hundreds or up to 1200 μm. Comparing PT and IT neurons, axonal end-points of IT neurons on average have larger SD of the coordinates than those of PT neurons for all the three coordinates, with the most prominent difference for *x* coordinate (Fig. 3A) (Welch’s *t*-test, *x*: *p*=2.5×10^−6^, *d*=0.85; *y*: *p*=0.019, *d*=0.42; *z*: *p*=0.0024, *d*=0.55). The same tendencies also appear when analysis is limited to neurons in MOp or MOs (Fig. 3B,C) (MOp, *x*: *p*=0.0076, *d*=0.86; *y*: *p*=0.014, *d*=0.81; *z*: *p*=0.10, *d*=0.53; and MOs, *x*: *p*=0.0020, *d*=0.79; *y*: *p*=0.028, *d*=0.54; *z*: *p*=0.029, *d*=0.59). It is therefore suggested that the ipsilateral intra-striatal axonal end-points of IT neurons are on average more spatially extended than those of PT neurons, especially for the medial-lateral direction.

**Figure 3.**
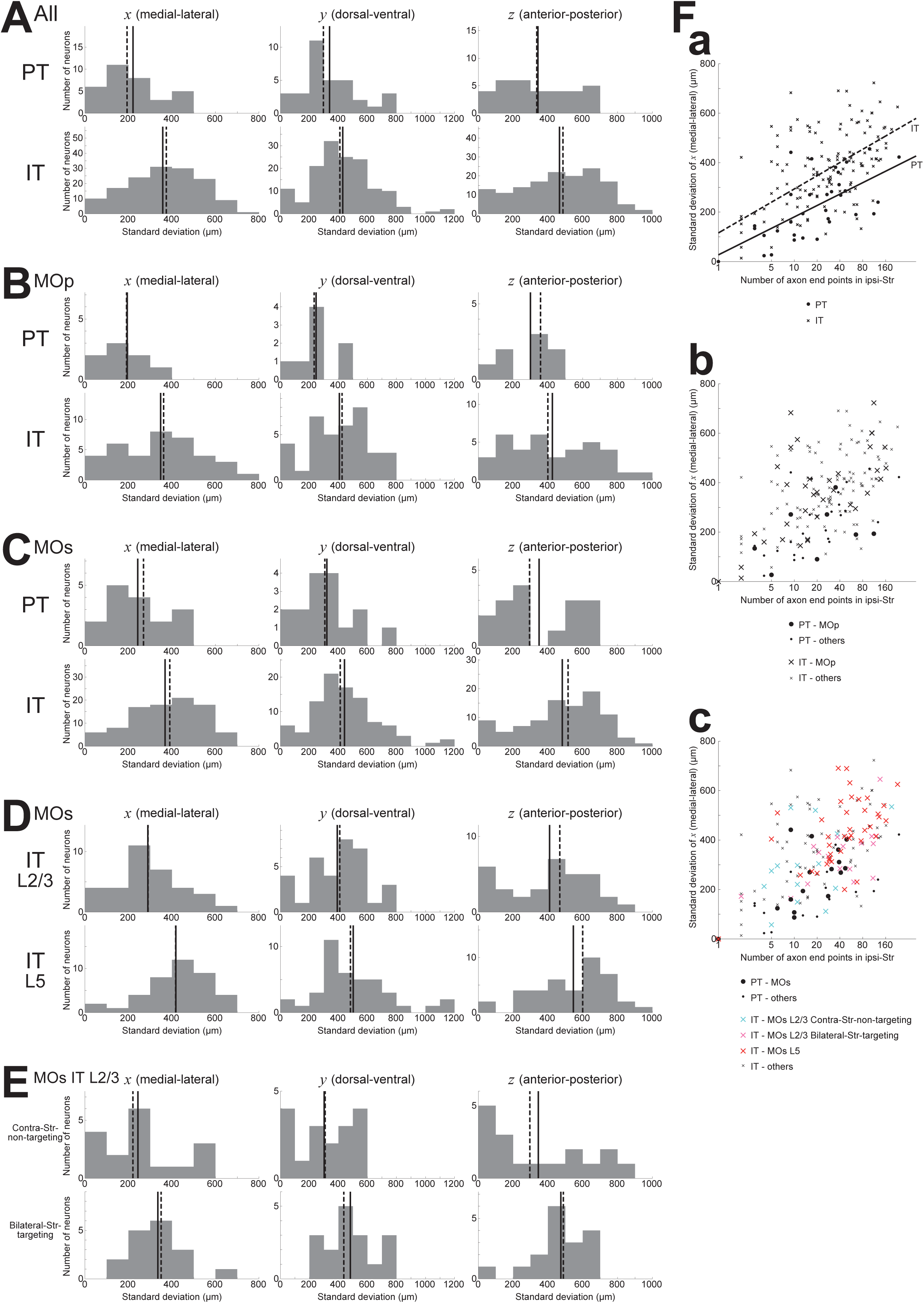
Distributions of the standard deviation (SD) of the spatial coordinates of ipsilateral intra-striatal axonal end-points. **(A)** The left, middle, and right panels show the distributions of the SD of *x, y*, and *z* coordinates (corresponding to the medial-lateral (or right-left), dorsal-ventral (or top-bottom), and anterior-posterior directions) of ipsilateral axonal end-points, respectively, for the striatum-targeting 33 PT (top panels) and 145 IT (bottom panels) neurons that have at least one ipsilateral end-point; i.e., IT neurons having no ipsilateral axonal end-point (16 neurons) are excluded from the histograms (the same is applied also to the following histograms). **(B**,**C)** Results for the neurons whose somata are located in the primary motor area (8 PT and 37 IT neurons (excluding 7 IT neurons w/o ipsi-point)) (B) or the secondary motor area (MOs) (16 PT and 94 IT neurons (excluding 7 IT neurons w/o ipsi-point)) (C). **(D)** Results for MOs layer 2/3 IT neurons (34 neurons, excluding 1 w/o ipsi-point) and MOs layer 5 IT neurons (40 neurons, excluding 3 w/o ipsi-point). **(E)** Results for MOs layer 2/3 IT neurons that do not target contra-Str (17 neurons) and those that target bilateral-Str (17 neurons). **(F)** Relationship between the logarithm of the number of ipsilateral axonal end-points (horizontal axis) and the SD of *x* coordinates of the end-points (vertical axis). **(a)** The circles and crosses indicate all the striatum-targeting PT and IT neurons that have at least one ipsilateral end-point, respectively. There is a positive correlation between the two variables in both PT neurons (*p*=6.4×10^−5^, *r*=0.64; *p*=3.9×10^−4^, *r*=0.59 when excluding a neuron with only one ipsilateral end point) and IT neurons (*p*=7.5×10^−17^, *r*=0.62; *p*=9.1×10^−10^, *r*=0.49 w/o neurons with one ipsi-point). The solid and dashed lines indicate the fitted lines of linear regression for PT and IT neurons, respectively (PT: intercept 27.0 (*p*=0.56), slope 66.7 (*p*=6.4×10^−5^); IT: intercept 115.8 (*p*=5.2×10^−5^), slope 77.2 (*p*<2×10^−16^)). **(b)** The neurons in MOp are indicated by large symbols. Positive correlation between the two variables exists for MOp IT neurons (*p*=2.9×10^−6^, *r*=0.69; *p*=1.6×10^−4^, *r*=0.60 w/o neurons with one ipsi-point), but not for MOp PT neurons (*p*=0.31; every neuron has >1 ipsi-points). **(c)** MOs PT neurons, MOs layer 2/3 IT contra-Str-non-targeting neurons, MOs layer 2/3 IT bilateral-Str-targeting neurons, and MOs layer 5 IT neurons are indicated by black large circles, light blue large crosses, pink large crosses, and red large crosses, respectively. Positive correlation between the two variables exists for MOs PT neurons (*p*=0.0059, *r*=0.66; *p*=0.065, *r*=0.49 w/o a neuron with one ipsi-point) and MOs entire IT neurons (*p*=9.4×10^−11^, *r*=0.61; *p*=2.5×10^−5^, *r*=0.43 w/o neurons with one ipsi-point).

We also analyzed if MOs IT neurons in layer 2/3 and those in layer 5 differ in the spatial distribution of ipsilateral intra-striatal axonal end-points. As shown in Fig. 3D, it turned out that the SD of the spatial coordinates of end-points is on average larger for MOs layer 5 IT neurons than for MOs layer 2/3 IT neurons in all the three directions, with the most prominent difference in the medial-lateral (*x*) direction (Welch’s *t*-test, *x*: *p*=7.8×10^−4^, *d*=0.82; *y*: *p*=0.045, *d*=0.47; *z*: *p*=0.013, *d*=0.60). The distributions differ also between MOs layer 5 IT neurons and MOs PT neurons (*x*: *p*=1.3×10^−4^, *d*=1.18; *y*: *p*=0.0054, *d*=0.75; *z*: *p*=0.0036, *d*=0.91) but not between MOs layer 2/3 IT neurons and MOs PT neurons (*x*: *p*=0.27; *y*: *p*=0.22; *z*: *p*=0.37).

As mentioned above, about a half of MOs layer 2/3 IT neurons (17/35) target bilateral striatum. The bilateral-Str-targeting neurons tend to have wider spatial distributions of ipsilateral end-points than contra-Str-non-targeting neurons (Fig. 3E) (Welch’s *t*-test, *x*: *p*=0.092, *d*=0.60; *y*: *p*=0.0074, *d*=0.99; *z*: *p*=0.12, *d*=0.55). The spatial distributions for the contra-Str-non-targeting neurons are narrower than layer 5 IT neurons (*x*: *p*=0.0019, *d*=1.06; *y*: *p*=0.0036, *d*=0.81; *z*: *p*=0.019, *d*=0.83), and comparable to PT neurons (*x*: *p*=0.99; *y*: *p*=0.79; *z*: *p*=0.96). By contrast, the spatial distributions for the bilateral-Str-targeting neurons are narrower in *x*-but comparable in *y*- and *z*-coordinates compared to layer 5 IT neurons (*x*: *p*=0.033, *d*=0.57; *y*: *p*=0.72; *z*: *p*=0.16), and wider than PT neurons (*x*: *p*=0.039, *d*=0.75; *y*: *p*=0.011, *d*=0.95; *z*: *p*=0.059, *d*=0.69).

We also examined the relationship between the logarithm of the number of ipsilateral end-points and the SD of their *x* coordinates. As shown in Fig. 3Fa, these two variables are positively correlated in both PT and IT neurons (see the legend for details). Moreover, results of linear regression of the SD of *x* coordinates against the logarithm of the number of end-points (PT: solid line; IT: dashed line; see the legend for details) indicate that IT neurons tend to have larger SD of *x* coordinates of end-points than PT neurons with comparable number of end-points. Also, the scatter plot distinguishing subpopulations of MOs IT and PT neurons (Fig. 3Fc) indicates that layer 2/3 contra-Str-non-targeting IT neurons (light-blue crosses) are distinct from layer 2/3 bilateral-Str-targeting (pink crosses) or layer 5 (red crosses) IT neurons and closer to PT neurons (black circles), in line with the results described in the previous paragraphs.

As a different measure of spatial extent of axonal end-points that unifies the three (i.e., *x, y*, and *z*) directions, we calculated the SD of the distances between the individual ipsilateral intra-striatal axonal end-points and the center of these end-points (i.e., the point whose coordinates are the means of the *x, y*, and *z* coordinates of individual end-points) for each neuron, excluding the neurons having no ipsilateral end-point (this is also applied to all the following analyses dealing with the SD of the distances). This SD of the distances turned out to be on average larger for IT neurons than for PT neurons (Fig. 4A) (Welch’s *t*-test, *p*=0.032, *d*=0.39), confirming that IT axonal end-points are spatially more extended than PT end-points. This relation also holds for MOp neurons only (*p*=0.036, *d*=0.73) or MOs neurons only (*p*=0.015, *d*=0.51). Distinguishing the layers of MOs IT neurons, the SD of the distances for layer 5 neurons is on average larger than that for layer 2/3 neurons (*p*=0.011, *d*=0.61), and the former is also larger than the value for MOs PT neurons (*p*=0.0012, *d*=0.86) whereas the latter is comparable to the value for MOs PT neurons (*p*=0.38). Further distinguishing bilateral-Str-targeting and contra-Str-non-targeting MOs layer 2/3 IT neurons, the SD of the distances for bilateral-Str-targeting neurons is larger than that for contra-Str-non-targeting neurons (*p*=0.011, *d*=0.92), and the former is also larger than the value for MOs PT neurons (*p*=0.021, *d*=0.85) and comparable to MOs layer 5 IT neurons (*p*=0.43) whereas the latter is comparable to the value for MOs PT neurons (*p*=0.50) and smaller than the value for MOs layer 5 IT neurons (*p*=0.0013, *d*=1.00).

**Figure 4.**
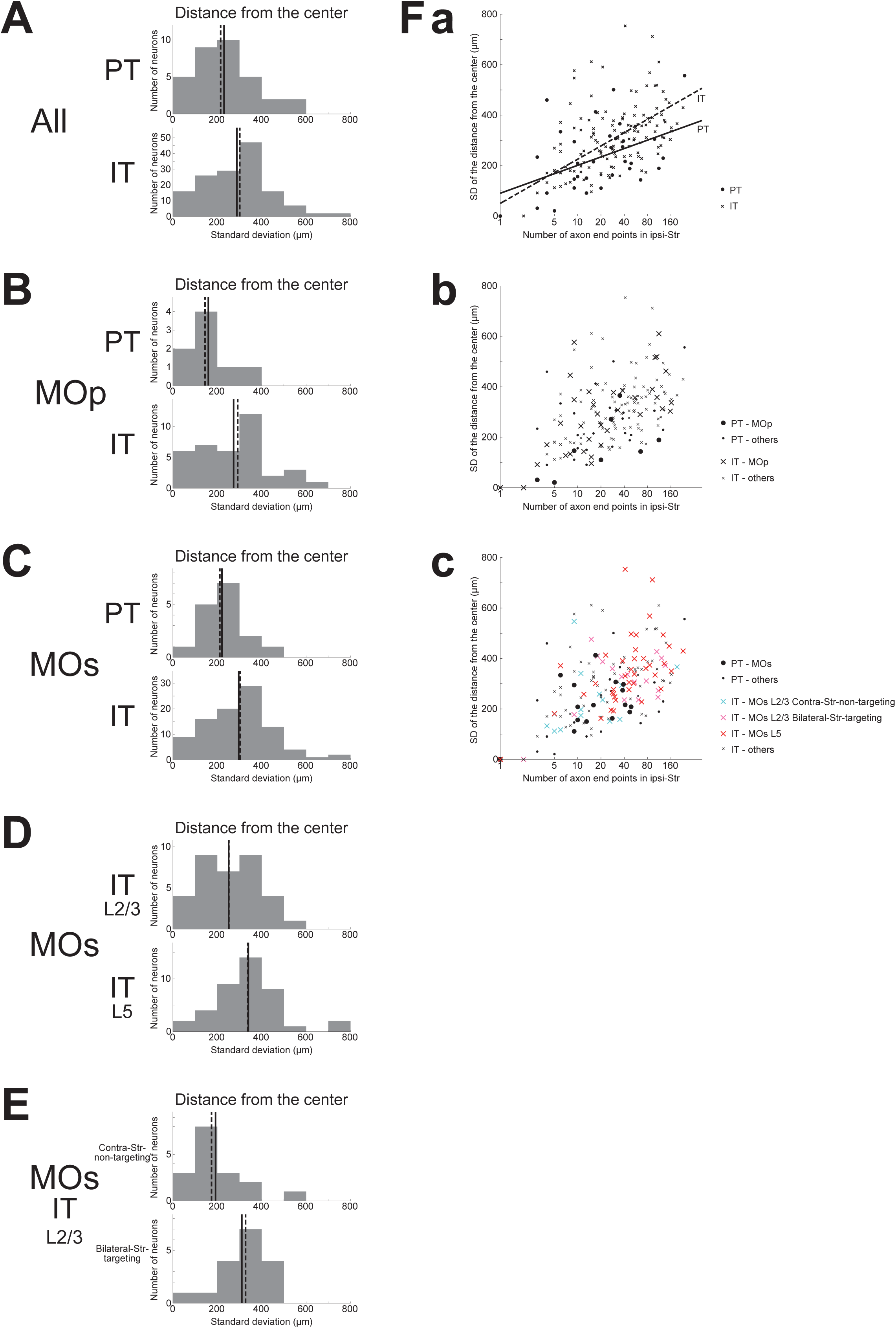
Distributions of the SD of the distances between the individual ipsilateral intra-striatal axonal end-points and the center of the end-points (i.e., the point whose coordinates are the means of the *x, y*, and *z* coordinates of individual end-points). **(A)** The top and bottom panels show the results for the striatum-targeting PT and IT neurons that have at least one ipsilateral end-point, respectively. IT neurons having no ipsilateral axonal end-point are excluded from the histograms, and the numbers of neurons included in, and excluded from, the histograms are the same as those in Fig. 3 (the same is applied also to the following histograms). **(B**,**C)** Results for the neurons whose somata are located in the primary motor area (MOp) (B) or the secondary motor area (MOs) (C). **(D)** Results for MOs layer 2/3 IT neurons and MOs layer 5 IT neurons. **(E)** Results for MOs layer 2/3 IT neurons that do not target contra-Str and those that target bilateral-Str. **(F)** Relationship between the logarithm of the number of ipsilateral axonal end-points (horizontal axis) and the SD of the distances of end-points from the center of end-points (vertical axis). **(a)** The circles and crosses indicate all the striatum-targeting PT and IT neurons that have at least one ipsilateral end-point, respectively. There is a positive correlation between the two variables in both PT neurons (*p*=0.0098, *r*=0.44; *p*=0.042, *r*=0.36 when excluding a neuron with only one ipsilateral end point) and IT neurons (*p*=6.4×10^−20^, *r*=0.67; *p*=1.8×10^−13^, *r*=0.57 w/o neurons with one ipsi-point). The solid and dashed lines indicate the resulting fitted lines of linear regression for PT and IT neurons, respectively (PT: intercept 90.3 (*p*=0.11), slope 48.1 (*p*=0.0098); IT: intercept 50.0 (*p*=0.041), slope 76.1 (*p*<2×10^−16^)). **(b)** The neurons in MOp are indicated by large symbols. Positive correlation between the two variables exists for MOp IT neurons (*p*=6.8×10^−7^, *r*=0.71; *p*=2.8×10^−5^, *r*=0.65 w/o neurons with one ipsi-point), and tends to exist for MOp PT neurons (*p*=0.10, *r*=0.62; every neuron has >1 ipsi-points). **(c)** MOs PT neurons, MOs layer 2/3 IT contra-Str-non-targeting neurons, MOs layer 2/3 IT bilateral-Str-targeting neurons, and MOs layer 5 IT neurons are indicated by black large circles, light blue large crosses, pink large crosses, and red large crosses, respectively. Positive correlation between the two variables exists for MOs entire IT neurons (*p*=1.6×10^−12^, *r*=0.65; *p*=8.4×10^−8^, *r*=0.53 w/o neurons with one ipsi-point), but hardly exists for MOs PT neurons (*p*=0.083, *r*=0.45; *p*=0.95, *r*=0.017 w/o a neuron with one ipsi-point).

We also examined the relationship between the logarithm of the number of end-points and the SD of the distances of end-points from the center of end-points. As shown in Fig. 4Fa, these two variables are positively correlated in both PT and IT neurons (see the legend for details). Results of linear regression of the SD of the distances against the logarithm of the number of end-points (PT: solid line; IT: dashed line; see the legend for details) indicate a somewhat steeper slope for IT neurons, but the difference between the PT and IT neurons is not drastic compared with the results of linear regression of the SD of *x* coordinates against the logarithm of the number of end-points (Fig. 3Fa). Meanwhile, the scatter plot distinguishing subpopulations of MOs IT and PT neurons (Fig. 4Fc) indicates that layer 2/3 contra-Str-non-targeting IT neurons (light-blue crosses) are distinct from layer 2/3 bilateral-Str-targeting (pink crosses) or layer 5 (red crosses) IT neurons and closer to PT neurons (black circles), in line with the results described in the previous paragraph and similarly to the results of linear regression of the SD of *x* coordinates against the logarithm of the number of end-points (Fig. 3Fa).

## Discussion

The present work addressed the long-standing issue, whether IT corticostriatal axons are morphologically more extensive than PT axons, by taking advantage of the recently developed public database of neuron morphology, in which we identified 33 and 161 striatum-targeting PT and IT neurons, respectively. Counting the number of intra-striatal axonal end-points, we have shown that there exists a large variety in the number of end-points across neurons in both neuron types. This variety seems in line with the suggested heterogeneity and existence of sub-types within each of PT and IT populations (Economo et al., 2018; Winnubst et al., 2019). More specifically, we found that, among MOs IT neurons, layer 5 neurons have a larger number of ipsilateral end-points than layer 2/3 neurons, and also bilateral-Str-targeting layer 2/3 neurons have a larger number of ipsilateral end-points than contra-Str-non-targeting layer 2/3 neurons. In contrast to these within-neuron-type differences, the entire IT and PT neurons turned out to be not drastically different in the number of ipsilateral end-points. This may be consistent with the previous study (Zheng and Wilson, 2002), which concluded that the once suggested difference between IT and PT axon morphology was spurious. Nonetheless, with the data of much increased number of neurons in the MouseLight database, we have shown that the proportion of neurons having >50 ipsilateral intra-striatal axonal end-points is larger in IT neurons than in PT neurons, although the difference is relatively small. Moreover, in MOs, layer 5 and bilateral-Str-targeting layer 2/3 IT neurons, but not contra-Str-non-targeting layer 2/3 IT neurons, have a larger number of ipsilateral end-points than PT neurons.

We have also examined the spatial extent of the distribution of ipsilateral axonal end-points, measured by the SD of the coordinates or of the distances from the center of end-points. With these measures, we have shown that IT ipsilateral axonal end-points on average have wider spatial distributions than PT end-points, with the difference along the medial-lateral axis most prominent. Distinguishing the subpopulations of MOs IT neurons, we have shown that layer 5 and bilateral-Str-targeting layer 2/3 IT neurons have a wider spatial distribution of ipsilateral axonal end-points than MOs PT neurons, whereas contra-Str-non-targeting layer 2/3 IT neurons are comparable to MOs PT neurons in these measures. Together with the abovementioned results for the number of axonal end-points, and considering that most layer 5 IT neurons target bilateral striatum (36/43 in MOs), it can be said, at least as for MOs, that bilateral-Str-targeting corticostriatal (CS) neurons generally have more extensive ipsilateral axons than contra-Str-non-targeting CS neurons in terms of the number and the spatial extent of end-points. The larger number and wider spatial extent of axonal end-points of bilateral-Str-targeting than contra-Str-non-targeting CS neurons suggests a possibility that former neurons affect a larger number of striatal neurons than the latter neurons, and functional significance of this would be interesting to explore.

As exemplified in this short article, the MouseLight database (Winnubst et al., 2019) is quite useful for testing the issues raised in previous anatomical and morphological studies with a smaller number of neurons. However, an important limitation is that information about synapses is not available in this database, in contrast to the previous studies that identified individual boutons (Zheng and Wilson, 2002) or even analyzed them by electron microscopy (Kincaid et al., 1998). Instead we analyzed the information about axonal end-points. However, it is not infrequent that a considerable portion of intra-striatal axons do not have any end-point (e.g., AA0182 and AA0011 PT neurons or AA0470 IT neuron in Fig. 5). This would cast doubt about whether the number of axonal end-points is well correlated with the number of synapses, and also about whether the spatial extent of axonal end-points well reflects the spatial extent of the entire intra-striatal axons. The latter issue is concerned also by the existence of cases where intra-striatal axons consist of multiple parts that are rather separate (e.g., AA0243 IT neuron in Fig. 5). These issues are expected to be complemented by future studies.

**Figure 5.**
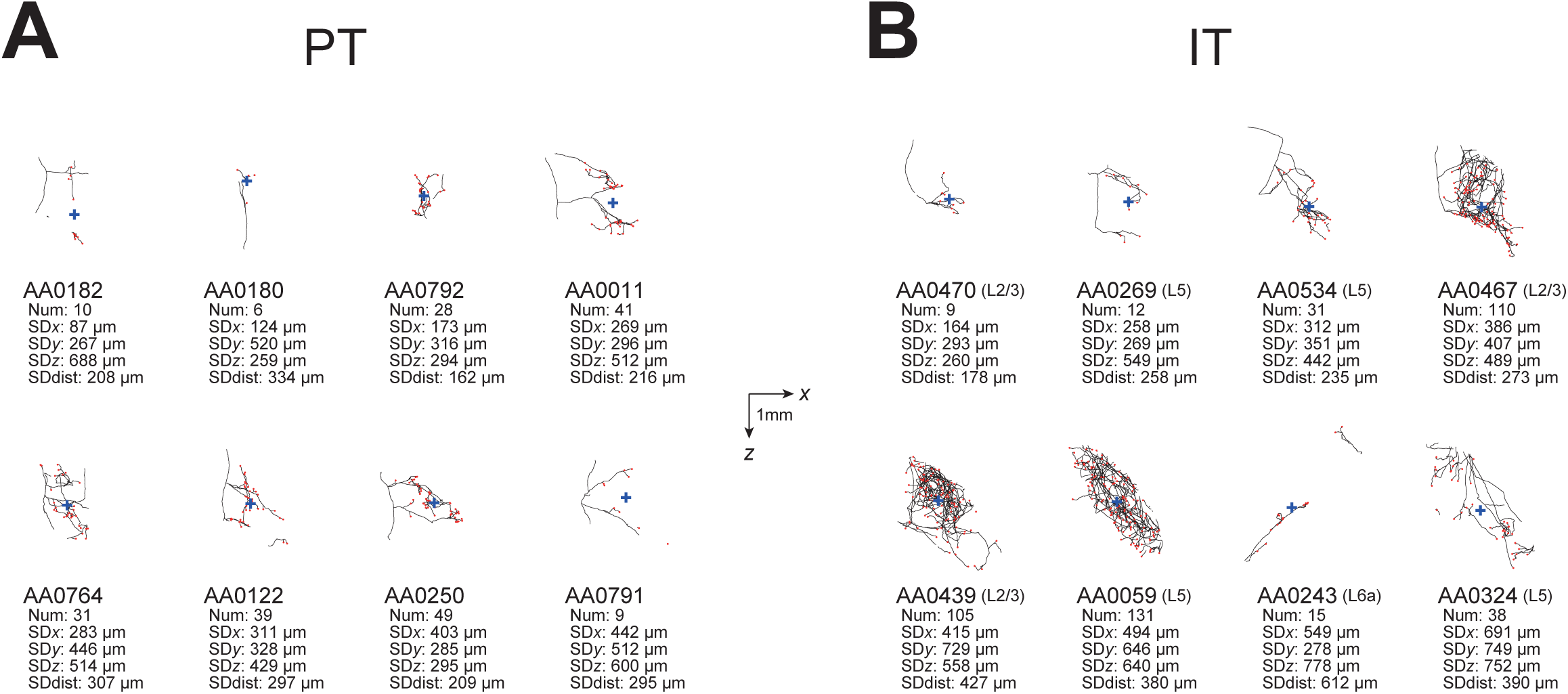
Examples of axonal arborizations of MOs PT and IT neurons in the striatum ipsilateral to the somata, drawn by using the data downloaded from the MouseLight database (http://ml-neuronbrowser.janelia.org/). Projections onto the horizontal section are drawn. The black lines and red points indicate the axons and axonal end-points, respectively, and the blue cross indicates the center of those end-points (i.e., the point whose coordinates are the means of the *x, y*, and *z* coordinates of individual end-points). The horizontal and vertical axes correspond to *x* (medial-lateral (or right-left)) and *z* (anterior-posterior) directions, respectively. The scale bars in the middle indicate 1 mm. The text under each panel describes the following information: ID number in the MouseLight database, with the layer of soma in the cases of IT neurons (B); Num: number of axonal end-points in the ipsilateral striatum; SD*x*,SD*y*,SD*z*: the standard deviation (SD) of *x, y*, and *z* coordinates of the ipsilateral intra-striatal end-points (in [μm]); and SDdist: SD of the distances of ipsilateral intra-striatal end-points from the center of those end-points (in [μm]). **(A)** Examples of PT neurons, whose SD of *x* coordinates of the ipsilateral intra-striatal axonal end-points (corresponding to the medial-lateral (or right-left) direction) are the 2, 4, 6, 8, 10, 12, 14, and 16-th from the smallest one among 16 MOs layer5 PT neurons (the order is from the top-left to top-right and then bottom-left to bottom-right). **(B)** Examples of IT neurons, whose SDs of *x* coordinates of the ipsilateral intra-striatal axonal end-points are the 10, 22, 34, 46, 58, 70, 82, and 94-th from the smallest one among 94 MOs IT neurons having at least one ipsilateral end-point (the order is from the top-left to top-right and then bottom-left to bottom-right).

## Acknowledgements

This work was supported by Grant-in-Aid for Scientific Research No. 15H05876 and 17H06311 of The Ministry of Education, Culture, Sports, Science and Technology in Japan and The Japan Society for the Promotion of Science to K. M. and Y. K., respectively.

## References

Arlotta P, Molyneaux BJ, Chen J, Inoue J, Kominami R, Macklis JD (2005) Neuronal subtype-specific genes that control corticospinal motor neuron development in vivo. Neuron 45:207–221.

Baker A, Kalmbach B, Morishima M, Kim J, Juavinett A, Li N, Dembrow N (2018) Specialized Subpopulations of Deep-Layer Pyramidal Neurons in the Neocortex: Bridging Cellular Properties to Functional Consequences. J Neurosci 38:5441–5455.

Brown SP, Hestrin S (2009) Intracortical circuits of pyramidal neurons reflect their long-range axonal targets. Nature 457:1133–1136.

Catsman-Berrevoets CE, Lemon RN, Verburgh CA, Bentivoglio M, Kuypers HG (1980) Absence of callosal collaterals derived from rat corticospinal neurons. A study using fluorescent retrograde tracing and electrophysiological techniques. Exp Brain Res 39:433–440.

Cowan RL, Wilson CJ (1994) Spontaneous firing patterns and axonal projections of single corticostriatal neurons in the rat medial agranular cortex. J Neurophysiol 71:17–32.

Economo MN, Winnubst J, Bas E, Ferreira TA, Chandrashekar J (2019) Single-neuron axonal reconstruction: The search for a wiring diagram of the brain. J Comp Neurol 527:2190–2199.

Economo MN, Clack NG, Lavis LD, Gerfen CR, Svoboda K, Myers EW, Chandrashekar J (2016) A platform for brain-wide imaging and reconstruction of individual neurons. Elife 5:e10566.

Economo MN, Viswanathan S, Tasic B, Bas E, Winnubst J, Menon V, Graybuck LT, Nguyen TN, Smith KA, Yao Z, Wang L, Gerfen CR, Chandrashekar J, Zeng H, Looger LL, Svoboda K (2018) Distinct descending motor cortex pathways and their roles in movement. Nature 563:79–84.

Gerfen CR, Paletzki R, Heintz N (2013) GENSAT BAC cre-recombinase driver lines to study the functional organization of cerebral cortical and basal ganglia circuits. Neuron 80:1368–1383.

Hooks BM, Papale AE, Paletzki RF, Feroze MW, Eastwood BS, Couey JJ, Winnubst J, Chandrashekar J, Gerfen CR (2018) Topographic precision in sensory and motor corticostriatal projections varies across cell type and cortical area. Nat Commun 9:3549.

Kincaid AE, Zheng T, Wilson CJ (1998) Connectivity and convergence of single corticostriatal axons. J Neurosci 18:4722–4731.

Kiritani T, Wickersham IR, Seung HS, Shepherd GM (2012) Hierarchical connectivity and connection-specific dynamics in the corticospinal-corticostriatal microcircuit in mouse motor cortex. J Neurosci 32:4992–5001.

Kuan L, Li Y, Lau C, Feng D, Bernard A, Sunkin SM, Zeng H, Dang C, Hawrylycz M, Ng L (2015) Neuroinformatics of the Allen Mouse Brain Connectivity Atlas. Methods 73:4–17.

Levesque M, Charara A, Gagnon S, Parent A, Deschenes M (1996) Corticostriatal projections from layer V cells in rat are collaterals of long-range corticofugal axons. Brain Res 709:311–315.

Miller R (1975) Distribution and properties of commissural and other neurons in cat sensorimotor cortex. J Comp Neurol 164:361–373.

Molyneaux BJ, Arlotta P, Fame RM, MacDonald JL, MacQuarrie KL, Macklis JD (2009) Novel subtype-specific genes identify distinct subpopulations of callosal projection neurons. J Neurosci 29:12343–12354.

Morishima M, Kawaguchi Y (2006) Recurrent connection patterns of corticostriatal pyramidal cells in frontal cortex. J Neurosci 26:4394–4405.

Morishima M, Morita K, Kubota Y, Kawaguchi Y (2011) Highly differentiated projection-specific cortical subnetworks. J Neurosci 31:10380–10391.

Parent M, Parent A (2006) Single-axon tracing study of corticostriatal projections arising from primary motor cortex in primates. J Comp Neurol 496:202–213.

Reiner A, Hart NM, Lei W, Deng Y (2010) Corticostriatal projection neurons - dichotomous types and dichotomous functions. Front Neuroanat 4:142.

Shepherd GM (2013) Corticostriatal connectivity and its role in disease. Nat Rev Neurosci 14:278–291.

Tasic B et al. (2018) Shared and distinct transcriptomic cell types across neocortical areas. Nature 563:72–78.

Wilson CJ (1986) Postsynaptic potentials evoked in spiny neostriatal projection neurons by stimulation of ipsilateral and contralateral neocortex. Brain Res 367:201–213.

Wilson CJ (1987) Morphology and synaptic connections of crossed corticostriatal neurons in the rat. J Comp Neurol 263:567–580.

Winnubst J, Bas E, Ferreira TA, Wu Z, Economo MN, Edson P, al. e (2019) Reconstruction of 1,000 projection neurons reveals new cell types and organization of long-range connectivity in the mouse brain. bioRxiv 537233.

Zheng T, Wilson CJ (2002) Corticostriatal combinatorics: the implications of corticostriatal axonal arborizations. J Neurophysiol 87:1007–1017.

